# NF-κB inhibitor alpha has a cross-variant role during SARS-CoV-2 infection in ACE2-overexpressing human airway organoids

**DOI:** 10.1101/2022.08.02.502100

**Authors:** Camille R. Simoneau, Pei-Yi Chen, Galen K. Xing, Mir M. Khalid, Nathan L. Meyers, Jennifer M. Hayashi, Taha Y. Taha, Kristoffer E. Leon, Tal Ashuach, Krystal A. Fontaine, Lauren Rodriguez, Bastian Joehnk, Keith Walcott, Sreelakshmi Vasudevan, Xiaohui Fang, Mazharul Maishan, Shawn Schultz, Jeroen Roose, Michael A. Matthay, Anita Sil, Mehrdad Arjomandi, Nir Yosef, Melanie Ott

## Abstract

As SARS-CoV-2 continues to spread worldwide, tractable primary airway cell models that accurately recapitulate the cell-intrinsic response to arising viral variants are needed. Here we describe an adult stem cell-derived human airway organoid model overexpressing the ACE2 receptor that supports robust viral replication while maintaining 3D architecture and cellular diversity of the airway epithelium. ACE2-OE organoids were infected with SARS-CoV-2 variants and subjected to single-cell RNA-sequencing. NF-κB inhibitor alpha was consistently upregulated in infected epithelial cells, and its mRNA expression positively correlated with infection levels. Confocal microscopy showed more IκBα expression in infected than bystander cells, but found concurrent nuclear translocation of NF-κB that IκBα usually prevents. Overexpressing a nondegradable IκBα mutant reduced NF-κB translocation and increased viral infection. These data demonstrate the functionality of ACE2-OE organoids in SARS-CoV-2 research and identify an incomplete NF-κB feedback loop as a rheostat of viral infection that may promote inflammation and severe disease.

## Introduction

Understanding how the SARS-CoV-2 virus replicates and causes damage in the upper and lower airways has been the focus of intense scientific research for the past 2 years. The virus replicates in airway epithelial cells, and its receptor ACE2 is expressed throughout the airways and lung, but less so from the airway to the alveolar space.^1^ Notably, replication of the Omicron variant appears to be confined to the upper airway tract, a finding that correlates with less severe disease.^2,3^

Removal and neutralization of potentially harmful substances from inhaled air are main functions of the airway epithelium, which consists of multiple cell types. 1) Basal cells are the progenitor cells that differentiate into the epithelial cell types. 2) Club cells secrete the immunomodulatory club cell secretory protein and also fulfill regenerative functions. 3) Goblet cells secrete mucins onto the internal surface of the respiratory tract, thereby forming a liquid layer (termed mucus) to protect the underlying epithelium. 4) Ciliated cells are located across the apical surface and facilitate the movement of mucus across the airway tract.^4^ In the lungs, alveolar epithelial cells (ATI and II cells) line the alveoli where gas exchange takes place.

The viral spike protein has been found in the airway epithelium and lung tissue of deceased COVID-19 patients.^5^ In *ex vivo* infections, ciliated cells were identified as natural targets of SARS-CoV-2.^6^ However, evidence of infection was also noted in secretory and basal cells, in both *in vivo* and *ex vivo* infections.^7^ This includes viral budding in cells with secretory vesicles in infected human bronchial epithelial cells;^8^ others found goblet cells infected in *in vitro* human airway epithelial cell cultures and, as a response to infection, showed increased mucus production.^9^

To recapitulate the complex cellularity of the airway epithelium and provide a physiological, yet workable model of the infection in humans, multiple stem-cell derived 3D organoid models have been applied to virology^7,10–12^. Organoids consists of multiple cell types grown in a 3D structure to mimic organ structure. They can be derived from embryonic stem cells (ESCs), pluripotent stem cells (iPSCs) or progenitor cells in adult tissues. SARS-CoV-2 replicates in iPSC- or ESC-derived human airway organoids (HAOs)^13,14^ and adult stem-cell HAOs that encompass both airway and alveolar cells^15,16^. In the latter model, donor lungs are digested and cultured in Matrigel and a medium that allows for the outgrowth of basal, club and goblet cells, which can then include ciliated cells with additional *ex vivo* differentiation steps.^4,17^

Differentiation, often performed at the air-liquid interface, produces a pseudostratified configuration that most closely resembles the *in vivo* situation, but is lengthy, requires large volumes of cells, and loses 3D structure. The undifferentiated organoids provide an intermediate model system that is more structured and heterogeneous than a cell line, yet less rigid than the fully differentiated model. These undifferentiated HAOs are tractable and easy to work with, while not being cancerously transformed and maintaining cellular diversity that resembles the airway epithelium. The undifferentiated organoid model has been used to study respiratory syncytial virus^17^ and SARS-CoV-2^6,18,19^. However, work with the latter is plagued with variable, donor-dependent infection rates where most organoids carry less than 5% infected cells^18,19^, a major obstacle for further analysis (i.e., single-cell genomics).

Single-cell RNA sequencing (scRNA-seq) has provided important insight into the biology of SARS-CoV-2 and is particularly valuable in heterogenous cell systems where it effectively separates infected and bystander responses across multiple cell types. ScRNA-seq data were generated throughout the pandemic in both clinical and laboratory settings to determine cell-intrinsic responses to the virus, map COVID-19 changes in the lung, and understand the tissue milieu of responses.^20–22^ These experiments, for example, distinguished interferon-stimulated gene (ISG) signatures in COVID-19 patients with mild to moderate phenotypes from patients with severe COVID-19 illness.^20^ At the same time, advances in computational modeling for single-cell genomics permit an integration of samples from multiple conditions or cohorts, in a manner that allows direct comparisons of cellular compositions and gene expression programs (viral and host) ^23^.

SARS-CoV-2 is no longer a uniform strain as multiple variants have arisen, both globally and regionally. The D614G mutation arose early in 2020, and the beginning of 2021 was characterized by the rise and fall of additional common variants, such as Alpha and Beta, which spread globally and eventually became regionally dominant. Others, such as Epsilon, spread locally before being outcompeted by a new variant^24^. Delta dominated infections by late 2021, until it was overtaken by an ever-evolving cast of Omicron variants^25,26^. This series of variants offers unique opportunities to delineate common and specific traits of SARS-CoV-2 infection and to better treat or predict clinical disease severity.

To address these challenges and opportunities for studying SARS-CoV-2 infection, we generated and explored an adult stem cell-derived undifferentiated HAO model. Our model was genetically modified with ACE2 overexpression to overcome obstacles with low infection rates and long and variable differentiation times. We studied the effects of infection in this system using a genetically diverse set of variants, and applied scRNA-seq to understand the cell-intrinsic response to SARS-CoV-2 in this primary cell model. We identified *NFKBIA*, a critical regulator of the NF-κB pathway, as a universal gene that marks highly infected cells across all viral variants, underscoring the recognized link between inflammation and SARS-CoV-2 infection that dominates acute respiratory distress syndrome and severe disease in the lung.

## Results

HAOs were grown from digested uninfected donor lungs, according to a published protocol (Fig. 1A).^17^ As expected and confirmed by light sheet microscopy, the majority of cells (~80%) in one organoid expressed P63, the basal cell marker, but a minority (~20%) represented secretory cells, including MUC5AC^+^ goblet and CC10^+^ club cells, which aligns with published reports (Fig. 1B).^17^ In the absence of cell differentiation, very few FOXJ1^+^ ciliated cells were observed (not shown). Compared to primary lung epithelium, markers for basal cells (P63, KRT5) were statistically overrepresented in quantitative RT-PCR analysis of organoids, and those for ciliated cells (FOXJ1), goblet cells (MUC5AC), and club cells (CC10) were underrepresented (Fig. 1C). Expression of ACE2 mRNA was also higher in primary lung tissue than undifferentiated organoids, and no significant difference was noted in TMPRSS2 mRNA levels (Fig. 1D,E). No ACE2 protein was detected in the organoids by western blotting, which explains their poor infection efficiency in previous studies (Fig. 1F,G). ^18^

**Figure 1:**
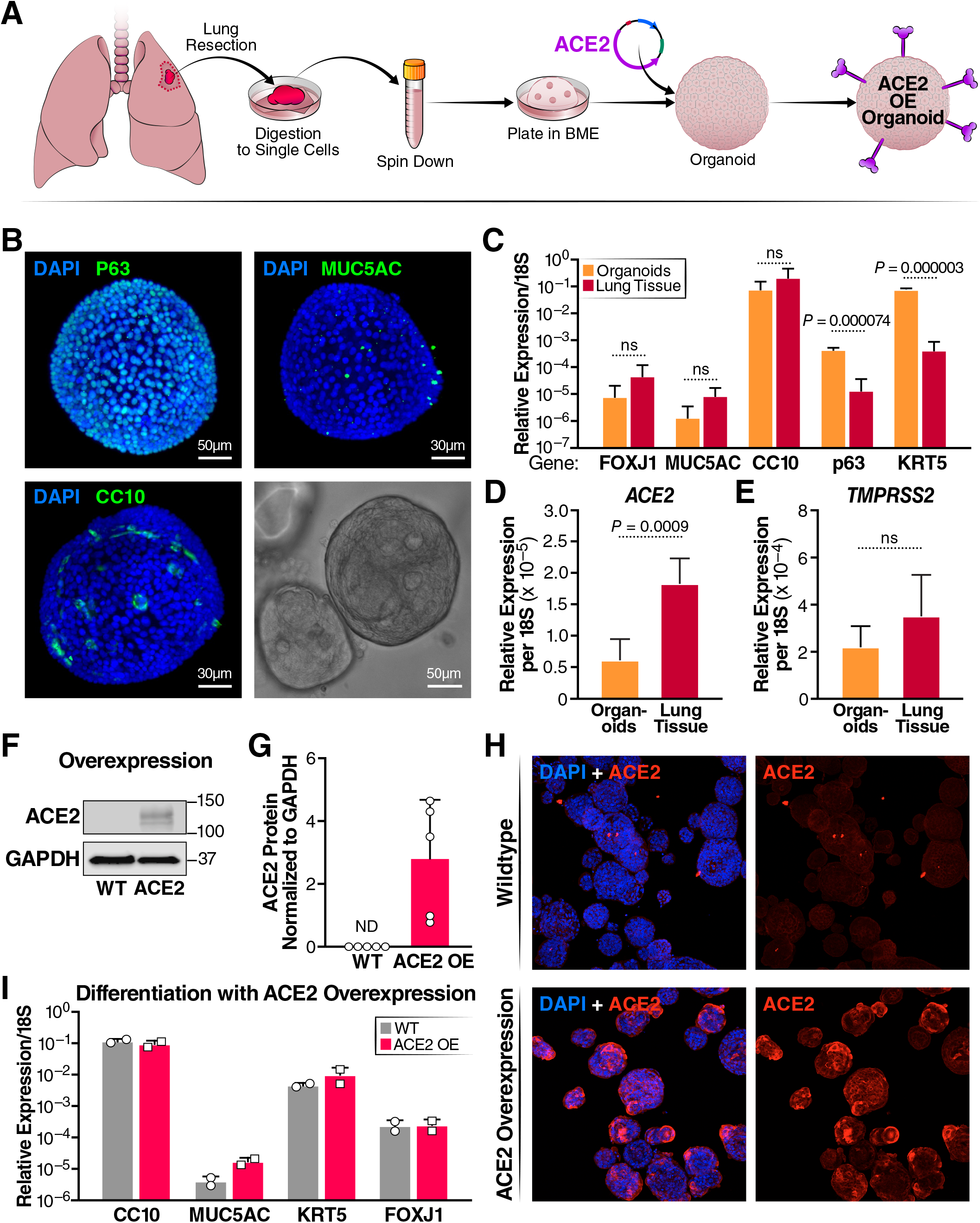
Characterization of human airway organoids (HAOs) and overexpression of ACE2. (A) Schematic detailing the establishment of organoid culture and ACE2 overexpression. (B) Light sheet microscopy of HAOs with staining for the cell markers CC10 (club cells) MUC5AC (goblet cells) and p63 (basal cells) with a brightfield image of a single organoid in the lower right-hand corner. (C) RT-qPCR comparison of cell-marker gene expression between digested primary lung tissue and organoids, (N = 3, organoids, N =4, input, +/−SD shown). (D) RT-qPCR comparing ACE2 expression level between organoids and digested lung tissue. (E) RT-qPCR comparing TMPRSS2 expression level between organoids and digested lung tissue (N = 3, organoids, N =4, input, +/−SD shown). (F) Representative western blot showing ACE2 protein levels in organoids at baseline and with ACE2 overexpression. (G) Quantification of ACE2 protein levels in wildtype organoids and organoids overexpressing ACE2. (H) Confocal imaging comparing ACE2 expression levels in wildtype and ACE2-overexpressing organoids. (I) RT-qPCR comparison of cell-marker genes after differentiation at the air-liquid interface of wildtype and ACE2 overexpressing organoids (N = 2, +/−SD shown).

To overcome these issues, organoids were transduced with a lentiviral vector expressing the *ACE2* open reading frame. ACE2-overexpressing organoids (ACE2-OE) were selected and maintained robust ACE2 levels over at least nine culture passages as shown by western blotting (Fig 1F, G). This was also confirmed by confocal immunofluorescence microscopy where ACE2 protein expression was visible on the surface of most cells in the ACE2-OE organoids (Fig. 1H). ACE2 overexpression did not affect the ability of organoids to differentiate at the air-liquid interface into all four cell types similarly to wildtype (WT) organoids as assessed by RT-qPCR for epithelial cell markers (Fig. 1I).

Next, WT and ACE2-OE HAOs were infected with the SARS-CoV-2 WA1 ancestral strain at a multiplicity of infection (MOI) of 1. Culture supernatants were collected at 24 and 72 hours and subjected to plaque assays. ACE2 overexpression resulted in approximately two log higher infectious particle production at 24 hours and a log difference at 72 hours (Fig. 2A-C). As a second measure of active viral infection, infected organoids were subjected to confocal immunofluorescence microscopy after staining with an antibody against double-stranded (ds) RNA. This antibody specifically stains the RNA replication centers in infected cells containing positive-negative strand RNA hybrids.^27^ dsRNA^+^ cells in WT and ACE2-OE organoids had two-to threefold greater ACE2 expression than WT organoids (Fig. 2D,E). These results show a robust increase in infected cell numbers with ACE2 overexpression that may now allow downstream analysis of single cells.

**Figure 2:**
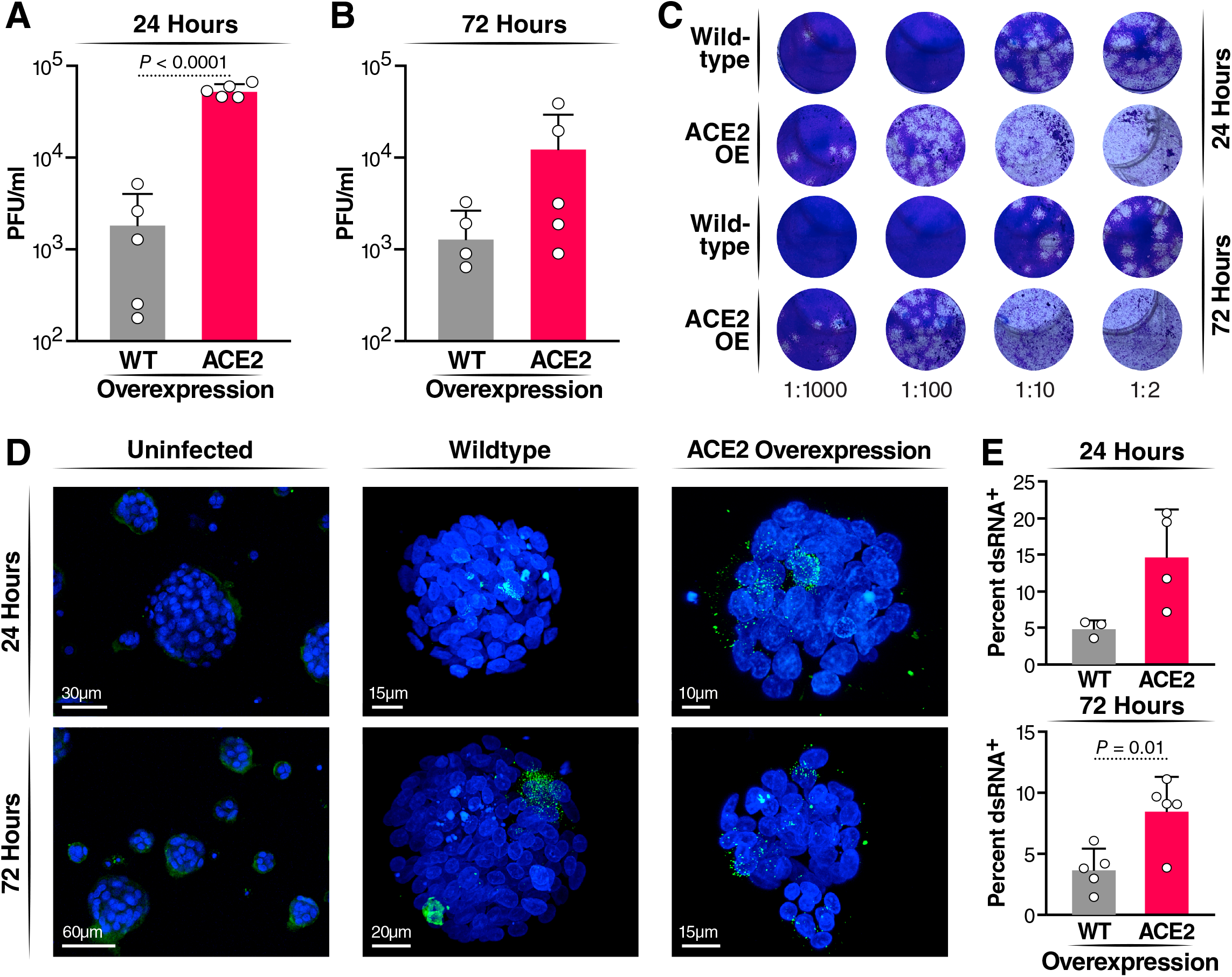
Infection of HAOs with SARS-CoV-2. Organoids were infected at an MOI of 1 for 2 hours, and then washed and fresh medium added. Organoids were incubated for the length of time indicated (A) Plaque assay comparing wildtype and ACE2 overexpressing organoids at 24 hours post-infection. (B) Plaque assay comparing wildtype and ACE2 overexpressing organoids at 72 hours post-infection. (C) Representative images of plaque assays performed on supernatant from SARS-CoV-2 infection of wildtype and ACE2 overexpressing organoids. (D) Representative images of infected HAOs at 24 and 72 hours stained for dsRNA (green). (E) Quantification of percent of total DAPI+ cells expression dsRNA across three experiments at 24 and 72 hours.

To test this, WT and ACE2-OE, organoids were subjected to 10X scRNA-seq after infection with the SARS-CoV-2 WA1 strain. Raw sequencing data were aligned to the human genome with the SARS-CoV-2 genome appended, run through scVI ^28,23^, and represented in a joint (batch-corrected) low dimensional space (see Methods). In that space, the WT infected (WT_I), ACE2 uninfected (ACE2_U) and WT uninfected (WT_U) conditions clustered together, and a large subset of the ACE2 infected (ACE2_I) cells clustered separately (Fig. 3A, B).

**Figure 3:**
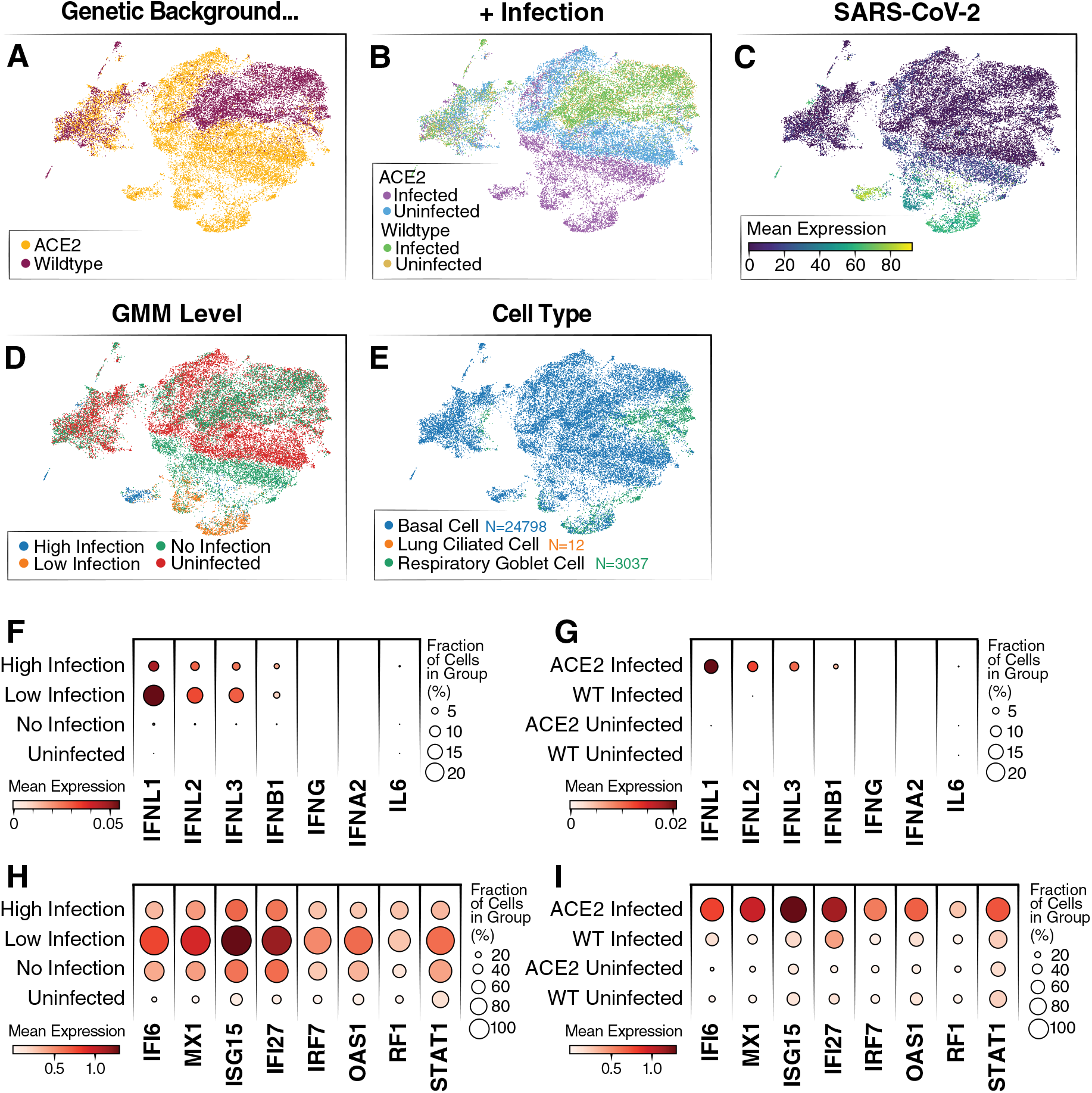
Single-cell RNA-sequencing of WT and ACE2 OE organoids infected with SARS-CoV-2. (A) UMAP colored by overexpression condition of organoids. (B) UMAP colored by overexpression condition and infection condition. (C) UMAP colored by log expression of SARS-CoV-2 RNA. (D) UMAP colored by GMM infection classification. (E) UMAP colored by cell type identification. (F) Dot plot showing levels of interferon genes and IL-6 RNA grouped by infection level. (G) Dot plot showing levels of interferon genes and IL-6 grouped by overexpression condition. (H) Dot plot showing levels of top differentially expressed ISGs grouped by infection level. ISGs were selected from the list of top 50 differentially expressed genes between the high-infected and uninfected cells. (I) Dot plot showing levels of top differentially expressed ISGs grouped by overexpression background.

To further validate the presence of cell types in the organoids identified by microscopy, the Lung Cell Atlas^29^ was used as a reference to assign cell types to our scRNA-seq data. In agreement with our previous results (Fig. 1), most cells were basal cells, and the remainder were respiratory goblet cells-a secretory cell type present in the lungs (Fig. 3E). Only a small number were identified as ciliated cells (Fig. 3E). Furthermore, this annotation revealed no significant difference in proportion of each cell type when ACE2 and WT organoids were compared, indicating that ACE2 overexpression does not change the cellular composition of organoids (Supp. Fig. 1A).

To further confirm that overexpression of *ACE2* did not change the biology of the organoids, a differential gene expression analysis was conducted between uninfected WT and ACE2-OE samples (Supp. Fig. 1B). *ACE2* was identified as the highest differentially expressed gene, and only four other genes (*C20orf85, BPIFA1, TMEM190, UTP14C*) were differentially expressed with a q-value less than 0.01. Considering SARS-CoV-2 RNA, we found expression primarily in cells from the ACE2_I condition (Fig. 3C, Supp. Fig1C). These results are consistent with previous data (Fig. 2), showing that SARS-CoV-2 infection levels are higher in the ACE2 condition than the WT. They further suggest that the difference in infection levels between the organoids is most likely due to ACE2 overexpression and not altered abundance of other receptors or cofactors.

Considering only cells from the infected conditions (ACE2_I, WT_I), we observed a broad variation in the abundance of viral products, presumably reflecting different levels of infection. To distinguish cells that had productive SARS-CoV-2 infections from those that had little to no infection, we fitted a Gaussian Mixture Model to classify cells from the infected samples into high, low and no infection groups (accounting for possible ambient viral product; see Methods and Fig. 3D). As expected, most of successfully infected cells (low and high infection groups in our model) came from the ACE2_I condition. Conversely, a subset of ACE2_I cells from the “no infection” group of cells clustered together with cells from the uninfected samples (ACE2_U, WT_U), suggesting that some cells in the infected organoid were not only uninfected but also not influenced by the infection of other cells.

To determine whether the successfully infected cells were detecting the viral replication, we assessed their levels of interferon gene expression. We found no expression of IFN-α, -β, or -γ in our scRNA-seq data. This result is similar to data published from COVID-19 patient samples in which IFN-λ is the primary response.^30^ Interestingly, while we found IFN-λ expression only in our infected samples, the highest amounts were observed in cells with low levels of infection, with substantially lower expression in cells from the no-infection or high-infection groups (Fig. 3F-G). While these data help to demonstrate that these organoids recapitulate the response of lungs during infection, they also suggest that the anti-viral IFN-λ response is only active in cells with low-level infections and shrinks as the amounts of intracellular viral products increase.

Unlike the ACE2_I sample, we found comparably little expression of interferon genes in the WT_I sample (Fig. 3G). To further interrogate this, we looked at the levels of interferon-stimulated genes (ISGs) in the different groups of cells. Consistently, we found that ISGs were primarily expressed in cells with low levels of viral RNA and not in the other infection groups or in the WT cells (Fig. 3H-I). Finally, for an unbiased analysis, we examined the differential expression in infected and uninfected samples in each of our genetic backgrounds (ACE2 OE and WT). Using gene set enrichment analysis, we found strong enrichment for interferon and anti-viral response program in ACE2 case, and much less so in the WT sample (Supp Fig. 2A-B).

Our results suggest that the lack of viral replication in WT cells is not due to an immune response, but more likely is a result of limited viral entry and replication. It therefore supports the notion that the WT system is not a good model for studying cellular response to SARS-CoV-2 infection, as the infection level is too low to stimulate an immune response. Conversely, due to the higher levels of replication in the ACE2 organoids, a measurable cellular response in those cells, and no difference in baseline biology between the ACE2 and WT organoids, we chose the ACE2 OE condition for follow-up scRNA-seq experiments.

Next, we compared transcriptome responses at the single-cell level in a second set of ACE2-OE organoids infected with various viral variants. Besides WA1 (using similar conditions as above), we tested Alpha (B.1.1.7), two separate isolates of the Beta variant (B.1.351), and the California-resident Epsilon variant (B.1.429) to capture the emerging diversity in viral sequences at the time of our study. Irrespective of cell type, the cells from the infected organoids were well mixed, and most uninfected cells clustered separately (Fig. 4A), indicating no clear difference or outlier among the infections (Fig. 4B). Basal cells remained the overwhelming cell type in this experiment, with a smaller representation of goblet cells (Fig. 4C). We again use our GMM approach for classifying the level of infection within each cell (Fig. 4D). As in Figure 3, we saw activation of INF-λ and ISGs across all samples infected by the variants but not in the uninfected sample (Fig. 4E-F).

**Figure 4:**
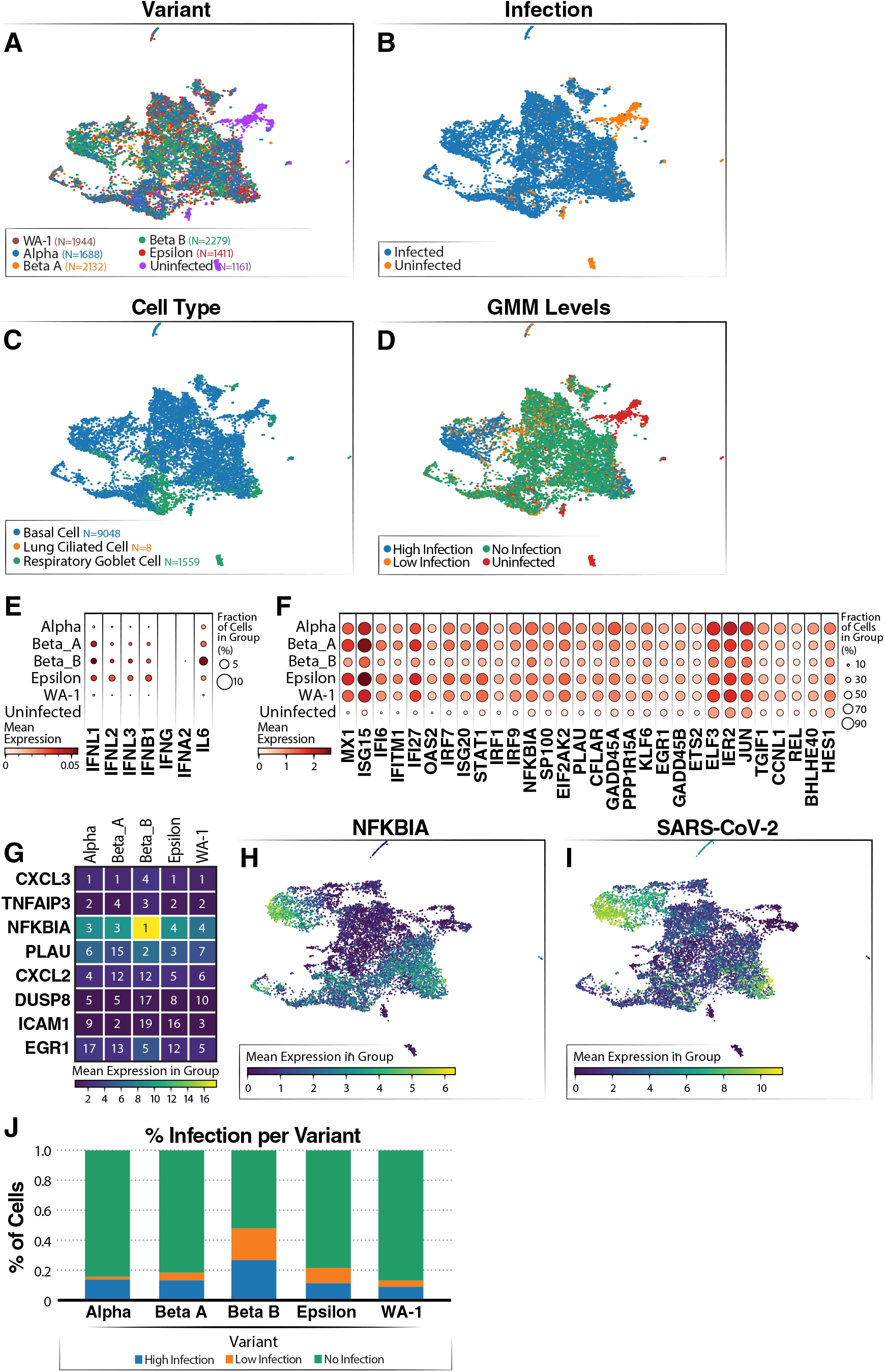
Single-cell RNA-sequencing of airway organoids infected with SARS-CoV-2 variants. (A) UMAP showing distribution of cells from each variant infection. (B) UMAP colored by infected and uninfected conditions. (C) UMAP colored by cell type (D) UMAP colored by SARS-CoV-2 infection level (via GMM classification). (E) Dot plot showing levels of interferon genes and IL-6 grouped by variant. (F) Dot plot showing levels of ISG expression grouped by variant. (G) Heatmap showing top five genes correlated to SARS-CoV-2 RNA level grouped by variant. The number in the box shows the ranking of the gene based on correlation for the variant and coloring shows the mean expression of the gene listed on the left. (H) UMAP colored by log expression of *NFKBIA* RNA. (I) UMAP colored by log expression of SARS-CoV-2 RNA. (J) Infection level proportions per variant.

In addition, all variants had similar infection levels with one of the Beta strains having a slightly higher level (Fig. 4J). However, a comparison of the replication dynamics over time of Beta B, Beta A and WA-1 showed no significant differences, suggesting that the higher levels of virus are due to variability and not to an intrinsic difference in these viruses (Supp. Fig. 3). Our results suggest that the different strains have comparable effects on our organoid system.

To determine which genes were most consistently upregulated with infection across the variants, we computed the correlation between gene expression and viral product (over cells) for each variant. The genes were then ranked by their level of correlation, separately for each variant (Fig. 4G; Table S1). Interestingly, we found a high level of consistency among the resulting lists of top ranked genes, with *CXCL3* being the top ranked in four out of the five variants, *TNFAIP3* in the top three in four of the variants, and *NFKBIA* in the top five in all variants. *NFKBIA* had the highest expression of the top-correlated genes, and had high expression in the high infection cluster (Fig. 4H). This cluster also had the highest levels of SARS-COV-2 viral products (Fig. 4I). *NFKBIA* is a quintessential NF-κB response gene and encodes the IκBa protein, which in turn dampens NF-κB activity by preventing its translocation into the nucleus.^31^ This usually generates a negative feedback loop where NF-κB activity is terminated or temporarily restrained by the upregulation of IκBα. The fact that *NFKBIA* is the top-upregulated gene in the analysis indicates strong activation of the NF-κB signaling pathway in infected cells across variants.

To investigate this finding in greater depth, we confirmed upregulation of the IκBα protein by confocal microscopy. For better resolution, we used A549 epithelial cells overexpressing ACE2 as they form a homogenous monolayer on the cover slips. Cells were infected with SARS-CoV-2 WA-1 and co-stained for dsRNA and IκBα. Cells were imaged, and the mean fluorescence intensities (MFI) of IκBα were compared in dsRNA^+^ and dsRNA^-^ cells. IκBα protein expression was enhanced in infected (dsRNA^+^) versus noninfected bystander (dsRNA^-^) cells, mirroring the induction of *NFKBIA* transcript levels (Fig. 5A). In a parallel experiment, we co-stained infected A549-ACE2 cells with antibodies against dsRNA and the p65/RELA subunit of NF-κB and quantified p65 nuclear localization. 75% of dsRNA^+^ cells showed p65 nuclear localization, but only a few dsRNA^-^ cells did (Fig. 5B, C). This confirms a strong activation of the NF-κB signaling pathway only in infected, and not in uninfected, cells, which explains the consequent upregulation of *NFKBIA* as a response gene. However, the resultant increase in IκBα activity appears insufficient to downregulate or terminate NF-κB signaling in cells infected with SARS-CoV-2, as the majority of infected cells localize p65 in the cell nucleus.

**Figure 5:**
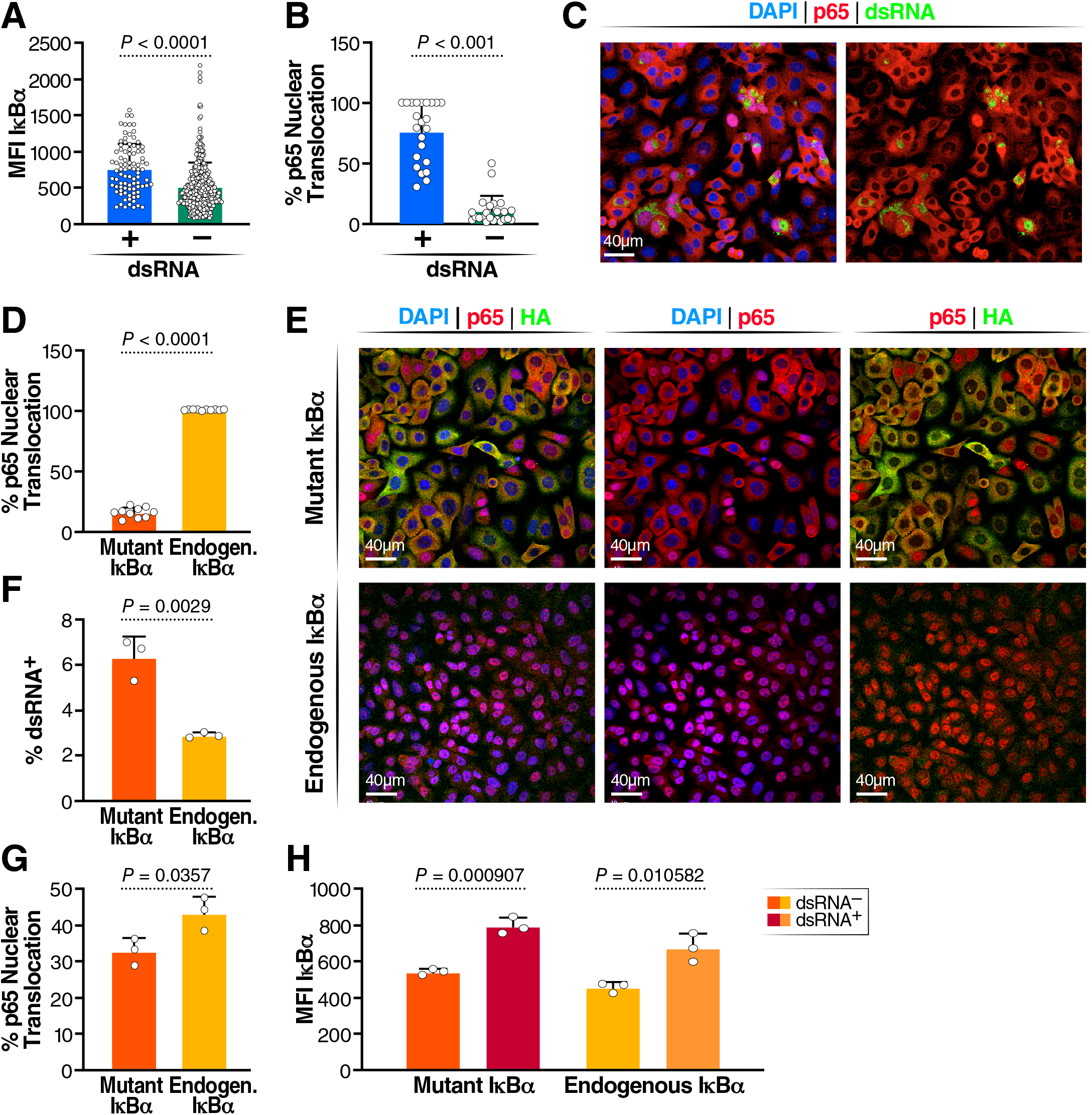
Role of IκBα in SARS-CoV-2 infection. (A) Quantification of p65 translocation in SARS-CoV-2 infected dsRNA+ and dsRNA-A549-ACE2 cells (n = 2) (B) Representative image of p65 translocation (C) Quantification if MFI of IκBα in dsRNA+ and dsRNA-cells (D) Representative image of IκBα and dsRNA staining in SARS-CoV-2 infected A549-ACE2 cells (E) Quantification of p65 translocation in cells expressing phosphorylation resistant IκBα after stimulation with TNFα and cells with functional IκBα. Three replicates were done, and a minimum of 5,000 cells were analyzed per condition; standard deviation is indicated (F) Representative images of DAPI, p65 and HA staining after TNFα stimulation (G) Quantification of dsRNA in mutant and non-mutant IκBα A549-ACE2 lines (G) Quantification of dsRNA in mutant and non-mutant IκBα A549-ACE2 lines. Three replicates were done, and a minimum of 5,000 cells were analyzed per condition; standard deviation is indicated (H) Quantification of p65 translocation in mutant and non-mutant IκBα A549-ACE2 lines (I) Quantification of IκBα MFI in mutant and non-mutant IκBα A549-ACE2 lines.

The IκBα protein can be phosphorylated by upstream kinases in a process that destabilizes the inhibitor and facilitates p65 nuclear translocation and transcriptional activity.^32^ To determine the role of IκBα in the infectious process, we stably overexpressed an IκBα construct in A549-ACE2 cells with the serine residues at positions 32 and 36 mutated to alanine to prevent phosphorylation.^33^ This generates a “super-inhibitor” that is unresponsive to cell signals and is highly active in retaining p65 in the cytoplasm, preventing its transcriptional activity in the nucleus.^33^ We confirmed the proper function of the mutated inhibitor by stimulating uninfected A549-ACE2 cells with TNFα, a strong activator of the NF-κB pathway through activation of relevant upstream kinases, leading to phosphorylation and degradation of IκBα and consequent nuclear NF-κB translocation.^34^ In the parent cell line with endogenous IκBα, 100% of cells had nuclear translocation of p65 in response to TNFα (Fig. 5E). However, in cells expressing nondegradable IκBα, only 25% of cells showed p65 nuclear translocation, confirming that the super-inhibitor remained stable, keeping p65 in the cytoplasm. Co-staining of p65 and the HA-tagged IκBα mutant revealed that the remaining p65 translocation occurred in cells with low expression of the stable IκBα mutant (Fig 5 E).

When cells expressing the IκBα mutant were infected with SARS-CoV-2 WA1, infected cultures showed two-fold more dsRNA^+^ cells than the parent cell line (Fig. 5F). In support of this model, we found significantly less p65 translocation in cultures overexpressing the IκBα mutant than in the parent cell line (Fig. 5G). However, IκBα expression (measured as MFI) was increased in dsRNA^+^ cells in both mutant and WT IκBα-expressing cell lines, suggesting residual activation of the NF-κB signaling pathway despite the super-inhibitor and resulting upregulation of endogenous *NFKBIA* in infected cells in both conditions (Fig 5H).

## Discussion

Here we generated a genetically modified airway organoid system that overcomes natural variations in ACE2 levels and allows high infection efficiency without lengthy differentiation. We anticipate that the tripling of infection rates for SARS-CoV-2 and its variants will also be observed for other ACE2-utilizing coronaviruses, such as SARS-CoV and hCOV-NL63. The overall increase in infected cells allowed meaningful single-cell transcriptomics analysis with sufficient infected versus uninfected bystander cell counts recorded for computational analysis. We found infection in three cell types: stem-like basal cells and secretory goblet and club cells, with high viral replication associated with infection level. Transcriptional profiling in all three cell types yielded common transcriptional changes as indicated by clustering according to infection status by different viral variants. *NFKBIA* stood out as a gene of interest in controlling SARS-CoV-2 infection across variants and a highly induced gene whose mRNA levels positively correlated with viral RNA levels. We confirmed that the encoded IκBα protein is expressed at increased levels in infected cells and that, despite this upregulation, NF-κB was found in the cell nucleus in infected, but not uninfected, cells as a sign of continuous activation of NF-κB signaling (Fig. 6).

**Figure 6:**
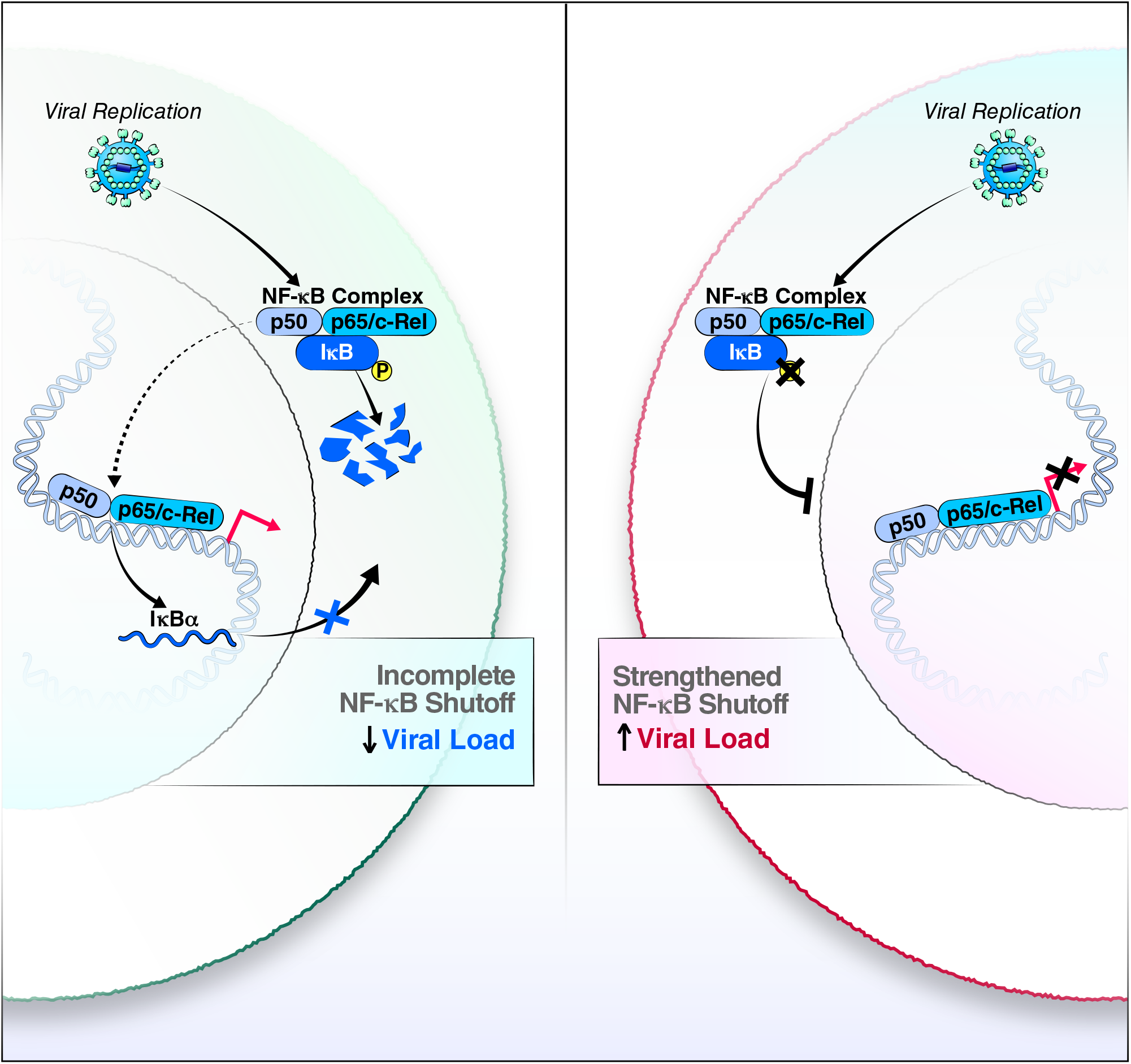
Model of incomplete NF-κB feedback loop in SARS-CoV-2-infected airways cells. While cells with endogenous IκBα maintain active NF-κB transcription factors in the cell nucleus and viral replication is attentuated (left), the negative feedback loop is strengthened with overexpression of an IκBα mutant, a process that enhances production of infectious SARS-CoV-2.

A similar incomplete feedback loop in NF-κB control was previously seen for respiratory syncytial virus infection^35,37,38^. However, the cause and benefit of this phenotype for viral infections remain unknown. It could represent an ongoing ‘arms race’ between the host and virus, where the host upregulates antiviral NF-κB signaling, but continuous viral presence overwhelms the feedback loop despite high rates of *NFKBIA* transcript and IκBα protein production through continuous degradation of the IκBα inhibitor. These data are consistent with published reports showing high levels of expression of NF-κB signaling-related genes in SARS-CoV-2-infected cells^35^. The overexpression of a mutant IκBα protein that cannot be phosphorylated and consequently degraded and the resulting two log increase in virus production support this model and identify IκBα as a proviral factor (Fig. 6). This is corroborated by studies by Sunshine et al., who identified *NFKBIA* in a Perturb-Seq study as a host protein supporting viral replication.^36^

It remains to be determined what the relevant upstream and downstream signals of this proviral phenotype are and whether the continuous presence of NF-κB in the cell nucleus provides benefits for the SARS-CoV-2 lifecycle. Our findings indicate that after overexpression of the mutant IκBα protein, nuclear p65 levels were lower in infected cells, supporting a model in which IκBα promotes viral infection by constraining, albeit not completely, the well-known antiviral activities of NF-κB. However, knockdown studies of p65 have pointed to a positive role of NF-κB signaling in SARS-CoV-2 infection that needs further mechanistic exploration.^37^ The fact that nuclear p65 is a striking differentiator of infected from bystander cells in our confocal microscopy studies underscores a possible role of nuclear NF-κB in SARS-CoV-2 infection.

While the causes of the observed phenotype remain a matter of speculation, the consequences are more predictable: a constitutively active NF-κB signaling pathway is likely to turn infected cells in the airway into inflammatory “factories” with overproduction of pro-inflammatory cytokines and chemokines that may activate the immune system and contribute to the cytokine storm that is characteristic of severe COVID-19.^38^ Indeed, we identify various TNFα-induced proteins, such as CXCL8 and other cytokines, as gene products correlated with increased infection (Table S1).

The organoid model presented here adds another tool to the studies of SARS-CoV-2 infection in primary airway cells. It overcomes the hurdle of donor-dependent variable infection rates^18^ and presents a tractable intermediate between cell line studies and fully differentiated primary cell models. It also offers insight into the transcriptional reprogramming occuring in progenitor cells of the airways in response to SARS-CoV-2 infection that may contribute to post-acute sequelae of viral infection^7,38,39^. We confirm that, in non-ACE2-OE organoids, infection of the three predominant cell types occurs, albeit at a much lower level than in ACE2-OE organoids. Our data showing that ACE2-OE organoids recapitulate transcriptional changes seen *in vivo*, such as the induction of IFN-λ, support the validity of the model.

As IκBα is a common factor in all variants tested, we envision universal application for our findings across variants. We speculate that expression of *NFKBIA* mRNA, IκBα protein or the degree of nuclear translocation of NF-κB may be markers of infection rates and possibly disease severity in infected individuals; these parameters may also distinguish more severe variants, such as Delta, from less pathogenic variants, such as Omicron. Caution is warranted when therapeutically interfering with NF-κB activity: inhibiting NF-κB signaling with drugs, such as bortezomib, a proteasome inhibitor that stabilizes the IκBα protein may enhance SARS-CoV-2 infection. In contrast, therapeutic interventions that destabilize the IκBα protein should block viral infection but may also further exacerbate pro-inflammatory cytokine production in infected cells. Future studies will test these and other potential clinical applications of our findings.

## Materials and Methods

### Cell lines

A549 cells were cultured in complete DMEM (DMEM with 10% FBS, 1% penicillin-streptomycin, 1% glutamine). A549-ACE2 cells were made as described^40^ and cultured in complete DMEM supplemented with 10 μg/mL blasticidin.

A549-ACE2-IκBα (SS32,36AA) were transduced with a lentiviral vector expressing IκBα (SS32,36AA) and selected in complete DMEM with 2 μg/mL of puromycin and 10 μg/mL of blasticidin. pLVX-EF1α-IκBα (SS32,36AA)-IRES-Puro was cloned by PCR amplification of HA-tagged IκBα (SS32,36AA) from pCMV4-3 HA/IκBα (SS32,36AA) and Gibson assembly, PCMV4-3 HA/IκBα (SS32,36AA) was a gift from Warner Greene (Addgene plasmid #24143) and pLVX-EF1α-IRES-Puro was purchased from Clontech Laboratories (plasmid #631988).

293T-HA-R-Spondin1-Fc cells were purchased from Trevigen (Catalogue Number 3710-001-K) and cultured according to the manufacturer’s protocol to generate conditioned medium of R-spondi-1. Briefly, cells were grown in selection growth media (DMEM with 10% FBS, 1% penicillin-streptomycin, 1% glutamine, and 100 mg/mL Zeocin) for >5 days until they were >90% confluent. Medium was replaced with organoid basal media (Advanced DMEM/F12 from Invitrogen supplemented with 1% penicillin-streptomycin, 1% Glutamax, and 10 mM HEPES). After 3 days, cell supernatant (i.e., R-spondin-1 conditioned medium) was collected, centrifuged at 3,000 x g for 15 min, filtered through a 0.22-µm filter, and frozen at −20°C in 10-mL aliquots. This process was repeated by adding fresh organoid basal medium to the cells and collecting supernatant after 4 days.

### Culture of human airway organoids

To generate HAOs, input cells were used from whole-lung lavages or from digestion of whole lungs. HAOs were generated from these cells as described ^17,41^. Briefly, single cells were suspended in 65% reduced growth factor BME2 (Basement Membrane Extract, Type 2, Trevigen, catalogue number 3533-001-02). From this mixture, 50-µL drops containing 1,000–40,000 cells were seeded in 24-well suspension culture plates (GreinerBio-one, catalogue number 662-102). Drops were incubated at 37°C for >20 min and solidified. After this, 500 µL of HAO medium was added to each well. HAO medium is organoid basal media (Advanced DMEM/F12 supplemented with 1% penicillin-streptomycin, 1% Glutamax, and 10 mM HEPES) supplemented with 10% (vol/vol) R-spondin1 conditioned medium, 1% B27 (Gibco), 25 ng/mL Noggin (Peprotech), 1.25 mM N-acetylcysteine (Sigma-Aldrich), 10 mM nicotinamide (Sigma-Aldrich), 5 nM Herefulin beta-1 (Peprotech), and 100 µg/mL of Primocin (InvivoGen). HAO medium was further supplemented with 5 µM Y-27632, 500 nM A83-01, 500 nM SB202190, 25 ng/mL of FGF-7, and 100 ng/mL of FGF-10 (all from Stem Cell Technologies), and HAO medium was replaced every 3–4 days.

After 14–21 days, organoids were passaged. For this, cold basal medium was used to collect organoids in 15-mL Falcon tubes and dissolved in BME2. Tubes were centrifuged at 250 x g for 5 min at 4 °C. Medium was aspirated, 10x TrypLE Select (Gibco) was added to the organoids, and the mixture was incubated at 37°C for 5–10 min. Organoids were further dissociated by pipette mixing and then diluted in cold basal medium. After another spin and medium aspiration, cells were mixed with BME2 and seeded into new drops. After this initial passage, organoids were passaged every 10–21 days. Stocks of early-passage (P1-P3) organoid lines were prepared by dissociating organoids, mixing them with recovery cell culture-freezing medium (Gibco), and freezing them by standard procedures. These samples could be thawed and immediately cultured in HAO medium.

### Creation of HAO-ACE2 lines

HAO-ACE2 lines stably expressing hACE2 were generated by lentiviral transduction with plasmid LV-ACE2^42^, and selected with 2 µg/mL of blasticidin, as described in a study from our group.^43^

### Differentiation of HAOs at the air-liquid interface

HAOs were cultured at the air-liquid interface (ALI) for differentiation. Organoids were collected in cold basal medium and dissociated as described above. After spin and medium aspiration, cells were resuspended in warm basal medium with 2% (v/v) BME2 and seeded onto pre-coated 2% BME2 in 6.5-mm insert of a 24-well plate Transwell Permeable Support (Costar). Cells were incubated for 1 hour at 37°C and then 500 µL of warm HAO medium was added to the bottom wells of the 24-well plate. Cells were grown to confluency for 1 week before the ALI was established by removal of apical side medium and substitution of basolateral medium for 1:1 (v/v) HAO medium and with Pneumacult ALI basal medium (Stem Cell Technologies, 05002). Basolateral medium was changed every 3–4 days, and mucus was washed from the apical side Cells were differentiated at the ALI for at least 1 month.

### Real-time quantitative PCR

The Qiagen RNEasy and Zymo DirectZol RNA MiniPrep were used for RNA extraction and isolation according to manufacturer’s instructions. Final RNA concentrations were measured with a NanoDrop ND-1000. Total RNA was reverse-transcribed using oligo(dT)_18_ primers (Thermo Scientific), random hexamers primers (Thermo Scientific), and AMV reverse-transcriptase (Promega). cDNA was diluted to 5 ng/µL. Gene expression was assayed by real-time quantitative PCR using Maxima SYBR Green qPCR Master Mix (Thermo Scientific) on a Biorad C1000 real-time PCR system. The SYBR Green qPCR reactions contained 10 μL of 2x SYBR Green Master Mix, 2 μL of diluted cDNA, and 8 pmol of both forward and reverse primers. The reactions were run using the following conditions: 50°C for 2 min, 95°C for 10 mins, followed by 40 cycles of 95°C for 5 secs and 60°C for 30 secs. Relative values for each transcript were normalized to 18S rRNA. Gene primers used are listed in Table S2. For every qPCR run, three technical replicates per sample were used for each gene.

### Western blots

Organoids were collected from culture plate with cold basal medium, washed, and centrifuged. The resulting cell pellet was re-suspended in 100 μL of RIPA buffer with HALT protease. Sample was incubated on ice for 30 min, before sonicating at 40% power for 10 sec (Sonic Dismembrator Model 500). Sonicated cells were spun down, and supernatants were collected as cell lysates.

Cell-lysate protein concentrations were determined by Bio-Rad DC Assay. For each sample, 16-20 μg of protein was loaded. Appropriate volume of cell lysate was lysed with Tris-glycine SDS sample buffer, heated at 95°C for 5 min. Lysate samples were immediately loaded onto 4–20% gradient gels (Bio-Rad, catalogue # 4561095) for SDS-PAGE for 90 min at 120V. Gel was wet-transferred onto a 0.45 μm nitrocellulose membrane for 1 hour at 100V. Blots were blocked in 5% milk in 1x TBST solution for 1 hour in room temperature with gentle rocking, then incubated with primary antibodies in 5% milk in 1x TBST overnight at 4°C with gentle rocking. Blot was washed three times with 1xTBST and incubated with goat anti-rabbit and goat anti-mouse IgG HRP-conjugated secondary antibodies diluted at 1:5000 with 5% milk in 1xTBST for 1 hour at room temperature with gentle shaking. After washing the blots three times with 1xTBST, blots were developed with Lumi-Light Western Blotting Substrate (Roche) for 5 min in the dark and visualized on a chemiluminescence imager (Bio-Rad ChemiDoc MP).

### Whole-mount organoid staining

Organoids were processed for imaging as described ^44^. Briefly, organoids were removed from BME2 with 3x cold PBS washes then fixed in 2–3% paraformaldehyde for 30 min on ice and washed 3x in PBS. Fixed organoid samples were stored at 4°C for up to 2 months.

For staining, organoids were blocked in PBS supplemented with 0.5% Triton X-100, 1% DMSO, 1% BSA, and 1% donkey or goat serum. Organoids were blocked for several hours at room temperature. Blocking solution was removed and replaced with blocking solution containing primary antibodies diluted 1:500 or 1:250. Organoids were incubated with primary antibodies for 24 hours at 4°C. After this, organoids were washed 3x in PBS and incubated with secondary antibodies diluted 1:250 in PBS at room temperature for several hours. Organoids were washed 3x in PBS and stained with Hoescht before visualization. Organoids were imaged on both Zeiss Axio Observer Z.1 and Zeiss Lightsheet Z.1 microscopes. Images were processed using a combination of the Zeiss software, ImageJ 1.51f, and Imaris 9.3. Primary antibodies we used are listed in Table S2.

### Confocal imaging of HAOs

HAOs were dissociated as described above. Organoids were resuspended in warm HAO medium with 2% BME2 and seeded onto glass microscopy chamber slides (Thermo Fisher, 154453) pre-coated with 1% (v/v) Geltrex (Gibco, A1413301). Organoids were allowed to embed in the Geltrex layer before aspirating the media and fixed in 4% paraformaldehyde for 30 min on ice and washed 3x in PBS.

For staining, organoids were permeabilized in 1% Triton X-100 for 10 min at room temperature. Organoids were then blocked in 100 µL of blocking solution (5% donkey or goat serum, 1% BSA, 0.1% cold fish skin gelatin, 0.1% Triton X-100, 0.05% Tween 20) for 2 hours with gentle rocking at room temperature. Organoids were then incubated in primary antibodies diluted in antibody solution (3% serum, 0.1% cold fish skin gelatin, 0.3% Triton X-100, 0.05% Tween 20) overnight at 4°C with gentle rocking. After, organoids were washed three times in 1x PBS with 0.1% BSA, and incubated with secondary antibodies conjugated with fluorophores, diluted 1:400 in antibody solution, for 2 hours at room temperature with gentle rocking while protected from light. Organoids were washed three times in 1x PBS with 0.1% BSA and stained with Hoescht33342 for 10 min at room temperature. Polyester gasket was removed from the glass slide, and rectangular glass slide was mounted onto the HAOs with Gold antifade reagent (Invitrogen, P36934).

Images were taken on the Olympus FV3000RS confocal microscope and processed with Imaris 9.3.

### Infection of HAOs with SARS-CoV-2

HAOs were dissociated and re-suspended to final density of 100,000 cells per 300 μL of HAO medium and seeded onto culture plates pre-coated with BME2 at 37°C for 1 hour. The cells were allowed to settle and embed onto BME2 coating for 20-60 min at 37°C before adding HAO medium. 24 hours later, the organoids were infected at an MOI of 1 for 2 hours, and then then the virus-containing medium was removed, the cells were washed, fresh medium was added, and the cells were incubated until post-infection processing.

### Propagation of SARS-CoV-2 variants

All SARS-CoV-2 variants were propagated on Vero-E6 cells expressing human TMPRSS2 in an aerosol biosafety level-3 lab (aBSL3) lab. Stocks were titered via plaque assay on Vero-E6 TMPRSS2 cells as described^45^ and sequenced to confirm no novel cell-culture mutations.

### Single-cell RNA sequencing

HAOs were infected with SARS-CoV-2 as described above for 24 hours. Organoids were removed from the plate by dissolving the geltrax with cold basal medium and spun down. A single-cell suspension was made by digesting the organoids with 10X TrypLE for 15 min and passing them through a 40-μM filter. Library prep was done according to the standard 10X protocol with 6,000 cells loaded into the 10X Chromium Controller.

Libraries were sequenced on a NovaSeq6000, and the RNA was aligned with Cell Ranger v.6.0.0 to the human GRCh38 reference (10X Genomics, STAR aligner with the complete SARS-CoV-2 genome (MN985325.1) appended.^46^

### Analysis of SARS-CoV-2 sequencing

After alignment, cells that had >20% Unique Molecular Identifiers (UMIs) coming from mitochondrial genes were removed. Cells were also removed if they had fewer than 200 UMIs or greater than 300,000 UMIs. After filtering, the single-cell RNA-seq dataset contained 38462 cells and 36611 genes.

### scRNA-seq analysis with scVI

We used the Seurat v3 method^47^ as implemented in Scanpy v.1.8.1^48^ to select the top 4000 highly variable genes from the dataset, excluding the SARS-CoV-2 genes and ACE2. Then we ran scVI^28^ as implemented in scvi-tools v.0.13.0^23^ with default parameters and with each sequencing round treated a batch covariate. This resulted in a single latent space for all the data. We visualized the data by running the Scanpy function scanpy.pp.neighbor, followed by scanpy.tl.umap for each round of sequencing independently on the scVI latent space.

### Cell-type annotation

We first used seven different automated cell-type annotation methods to get a consensus prediction using the Lung Cell Atlas as a reference dataset.^29^ The seven methods were: 1. OnClass.^49^ 2. SVM. ^50^ 3. Random Forest (as implemented in sklearn).^51^ 4. KNN after batch correction with BBKNN.^52^ 5. KNN after batch correction with Scanorama.^53^ 6. KNN trained on the Lung Cell Atlas^29^ scVI latent space^28^. 7. scANVI.^54^ The resulting prediction was the majority prediction over the seven methods. We also derived a consensus score, based on the number of methods that agreed with the consensus prediction. After manual inspection, we determined that the only cell types present in the dataset were either: “Basal Cells”, “Goblet Cells”, and “Lung Ciliated Cells”. For all cells with a consensus score less than 5 or were not predicted as a basal/goblet/ciliated cell, we reclassified with a KNN trained on cells with consensus score greater than 5 and predicted as a basal/goblet/ciliated cell.

### Correlation analysis

To identify the top genes correlated with increased SCV2 levels, we computed the Pearson correlation as implemented in SciPy v1.4.1 (scipy.stats.pearsonr). We calculated the correlation for each gene in each sample individually between the log-normalized gene counts as implemented in Scanpy^48^ v.1.8.1 (scanpy.pp.normalize_total(adata, target_sum=1e4) followed by sc.pp.log1p) with the sum of all SCV2 RNA transcripts per sample. Then for each sample, we filtered for genes expressed in at least 20% of cells in the sample.

### DE testing and GO analysis

We used DESeq2 (v1.34.0) to identify genes differentially expressed between WT HAOs and ACE2-overexpressed HAOs by pseudobulking the transcript counts of each sample.^55^ We used Scanpy v.1.8.1 to identify differentially expressed genes between cells in the infected vs uninfected conditions.^48^ Metascape (v 3.5) with default Express Analysis settings were used to identify enriched Gene Ontology Terms.^56^

### Classifying infection with Gaussian Mixture Models

To classify whether a cell was actually infected with SCV2 (not just exposed to the virus), we used a Gaussian Mixture Model (GMM) to classify whether the SCV2 mRNA UMIs were due to background viral mRNA or from replicating virus within a cell. We used the scikit-learn v.1.0.1 GMM implementation (sklearn.mixture.GaussianMixture) with default parameters. Bayesian information criterion (BIC)^57^ as implemented in scikit-learn was used for model selection; defined as BIC = − 2log(L) + log(N)d, where L is the maximum likelihood of the GMM, N is the number of samples, and d is the degrees of freedom. For each experimental condition, we tested the optimal GMM for two vs three components as well as whether a single model should be trained on the log (sum of viral mRNA per cell + 1) or a separate model for each individual log(viral mRNA counts + 1) per viral gene. When training a separate model individually, we trained a GMM for each viral gene, summed the posterior probability of each component for each model, and for each cell assigned the component with the greatest total posterior probability. The BIC was then calculated as the mean BIC for each individual model. For each experimental condition, we selected the model with the lowest BIC. In all the uninfected experimental conditions (cells not exposed to virus), the optimal GMM model was a two-component GMM trained on the log (sum of viral mRNA per cell +1), while the optimal model for all the infected experimental conditions (cells exposed to virus), was a three-component GMM trained on the log(viral mRNA counts + 1) for each viral gene. For the two-component GMM, we interpreted the component with a lower mean as cells with “No Infection” while the higher component was “Yes Infection”. It is worth noting that all cells in the uninfected condition were classified as “No Infection,” which tracks with it being a negative control. For the three-component GMM, we interpreted the component with the lowest mean as “No Infection”, the middle mean as “Low Infection”, and the highest mean as “High Infection”.

## Acknowledgements

The following reagents were obtained through BEI Resources, NIAID, NIH: SARS-Related Coronavirus 2, Isolate USA-WA1/2020, NR-52281, SARS-Related Coronavirus 2, Isolate hCoV-19/South Africa/KRISP-EC-K005321/2020, NR-54008, contributed by Alex Sigal and Tulio de Oliveira (Beta_A), SARS-Related Coronavirus 2, Isolate hCoV-19/South Africa/KRISP-K005325/2020, NR-54009, contributed by Alex Sigal and Tulio de Oliveira. (Beta_B).

We thank Veronica Fonseca for administrative support, Mauricio Montano for BSL-3 laboratory support, Sara Gardner and John Carroll for graphics design, Dr. Norma Neff and Dr. Amy Kistler for sequencing support, the California Department of Public Health, especially Dr. Mary Kate Morris and Dr. Carl Hansen, as well as Dr. Charles Chiu, Dr. Raul Andino and Dr. Miguel Garcia Knight for the Epsilon variant and Alpha variant. We thank Adam Gayoso for feedback about single cell data analysis. We thank members from the laboratories of Melanie Ott, Nir Yosef, and Aaron Streets for general feedback. We gratefully acknowledge support from the CZ Biohub, the Innovative Genomics Institute and the Pendleton Foundation (M.O.), and Administrative COVID-19 Supplement 3P01AI091580-09S1 (J.P.R) on the parent NIH/NIAID P01-AI091580 (Weiss). SV and MA efforts were funded by the California Tobacco-related Disease Research Program (28FT-0020 to SV and T29IR0715 to MA) and the Departments of Defense (W81XWH-20-1-0158 to MA) and Veterans Affairs (CXV-00125 to MA).

## Author Contributions

CRS, P-YC, and GKX contributed equally. CRS: Conceptualization, Validation, Formal Analysis, Investigation, Writing-Original Draft, Writing-Review and Edit; P-YC: Investigation, Validation, Writing-Original Draft, GKX: Software, formal analysis, Writing-Original Draft, Writing-Review and Edit; MMK, JMH, TYT, KEL, SV, XF, MM: Investigation; TA, SS: Software; NLM, KAF: Conceptualization, Investigation; LR, BJ, KW: Resources; JR, MAM, AS, SV, MA: Resources, Supervision; NY, MO: Conceptualization, Funding acquisition, Supervision, Writing-Review and Edit.

## Figure legends

**Supplemental Figure 1: Related to Figure 3. Similarities and differences in ACE2 OE and WT organoids** (A) Proportion of goblet and basal cells in the ACE2 OE and WT backgrounds. Small numbers of ciliated cells (<25) were removed. (B) DEseq2 differential expression results of ACE2 vs WT cells. Highlighted cells have a q>0.01 (C) GMM infection classification of cells in the infected condition, broken down by cell type and genetic background. (D) Values of GMM infection classification of cells in the infected condition. (E) Proportion of goblet and basal cells in the infected vs uninfected conditions. (F) Proportion of goblet and basal cells in each of the classified infection levels (from GMM) and the uninfected condition.

**Supplemental Figure 2: Related to Figure 3. Gene Ontology enrichment analysis**. Metascape analysis showing top functional processes of genes differentially upregulated in cells in the infected vs uninfected conditions in an (A) ACE2 overexpression and (B) WT background.

**Supplemental Figure 3: Replication dynamics of Beta variants** (A) Quantification of dsRNA+ cells in A549-ACE2 at the indicated time in Beta variants and WA-1. (N =3, MOI = 1).

**Supplemental Figure 4: Related to Figure 3: Expression of viral entry factors**. Violin plots of SCV2 entry factors between ACE2 OE and WT HAOs in the infected and uninfected conditions. Plots show log normalized expression of (A) ACE2, (B) BSG, (C) NRP1, (D) TMPRSS2, (E) TMPRSS4, and (F) TSPAN8.

---

**Supplementary Table 1:** Top 25 genes correlated with SCV2 levels per variant.

|  | Alpha | Beta_A | Beta_B | Epsilon | WA-1 |
| --- | --- | --- | --- | --- | --- |
| 0 | CXCL3 | CXCL3 | NFKBIA | CXCL3 | DRD5 |
| 1 | TNFAIP3 | ICAM1 | PLAU | TNFAIP3 | CD33 |
| 2 | NFKBIA | NFKBIA | TNFAIP3 | PLAU | AC103808.1 |
| 3 | CXCL2 | AC091544.4 | CXCL3 | NFKBIA | AL357873.1 |
| 4 | DUSP8 | TNFAIP3 | EGR1 | CXCL2 | LRRIQ4 |
| 5 | PLAU | UBASH3A | CXCL8 | CXCL8 | LINC02284 |
| 6 | CXCL8 | NRAP | AC127526.5 | KLHL5 | BEST3 |
| 7 | KLF6 | AC010624.3 | NCOA7 | DUSP8 | U91328.3 |
| 8 | ICAM1 | AC008115.4 | EDN1 | CXCL1 | AC084816.1 |
| 9 | CXCL1 | DUSP8 | HES1 | NFKBIZ | CXCL3 |
| 10 | KLHL5 | AL450270.1 | KLF6 | LCN2 | TNFAIP3 |
| 11 | EDN1 | CXCL1 | CXCL2 | EGR1 | ICAM1 |
| 12 | BIRC3 | CXCL8 | NFKBIZ | TM4SF1 | NFKBIA |
| 13 | NFKBIZ | EDN1 | JUN | KLF6 | EGR1 |
| 14 | STX19 | KLHL5 | CXCL1 | PIM3 | LINC01354 |
| 15 | CCN1 | NR1D1 | AP003498.2 | ICAM1 | AL139118.1 |
| 16 | EGR1 | NFKBIZ | DUSP8 | AC127526.5 | AP001001.1 |
| 17 | TMEM156 | CXCL2 | ARRDC3 | AP003498.2 | RAB40AL |
| 18 | PIM3 | BHLHE41 | ICAM1 | AC055720.1 | AP006261.1 |
| 19 | AC127526.5 | EGR1 | TM4SF1 | AC010255.1 | AC032044.1 |
| 20 | IL6 | BIRC3 | EFNA1 | LINC01814 | NXPH3 |
| 21 | IL32 | PLAU | ZC3H12A | AC127537.1 | MYRFL |
| 22 | CD83 | KLF6 | AL627171.2 | EDN1 | AC022509.2 |
| 23 | NCOA7 | HIST1H2BD | KLHL5 | AC005062.1 | TNF |
| 24 | DUSP10 | IPCEF1 | MT-ND3 | SFRP4 | AC021483.2 |

**Supplementary Table 2:**
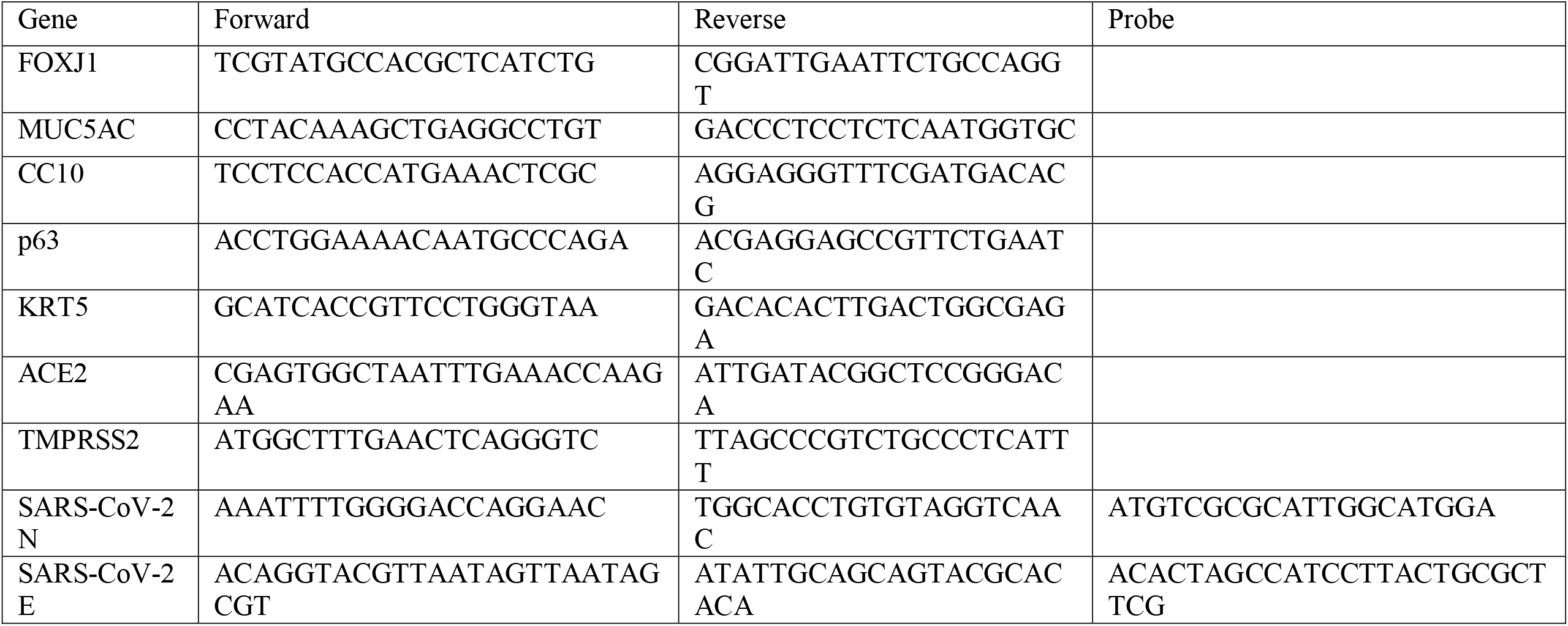
qPCR primers.

