## Supplemental Figure 1-4 for "NF-κB inhibitor alpha has a cross-variant role during SARS-CoV-2 infection in ACE2-overexpressing human airway organoids"

### Supplementary Figure 1

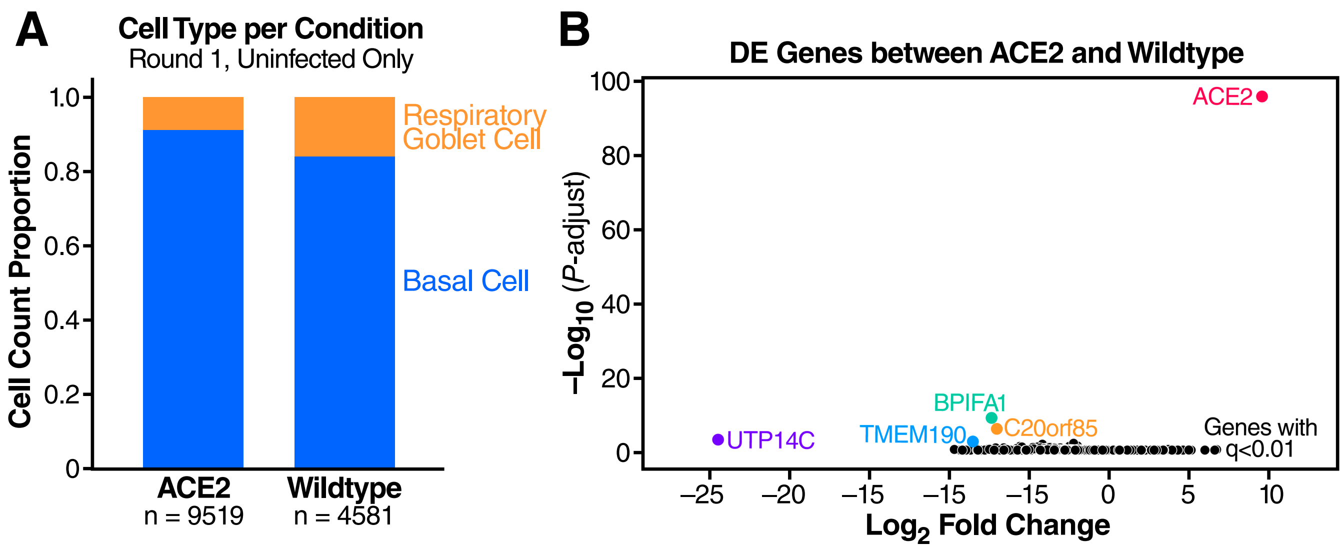


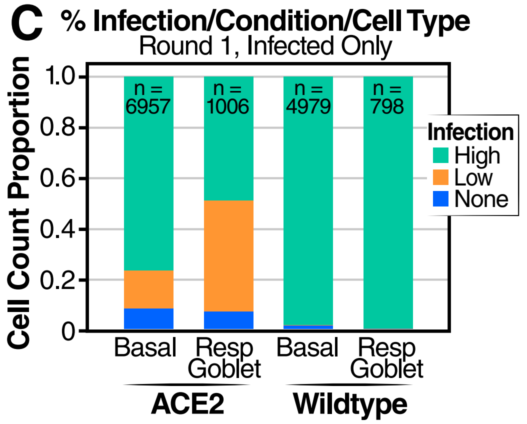


**D**

| | | Infection GMM Annotated | | |
|---|---|---|---|---|
| | | **High** | **Low** | **None** |
| **ACE2** | Basal | 8.24% n=573 | 15.01% n=1044 | 76.76% n=5340 |
| | Resp. goblet | 7.06% n=71 | 43.94% n=442 | 49.01% n=148 |
| **WT** | Basal | 1.47% n=73 | 0.04% n=2 | 98.49% n=4904 |
| | Resp. goblet | 0.25% n=2 | 0% n=0 | 99.75% n=796 |

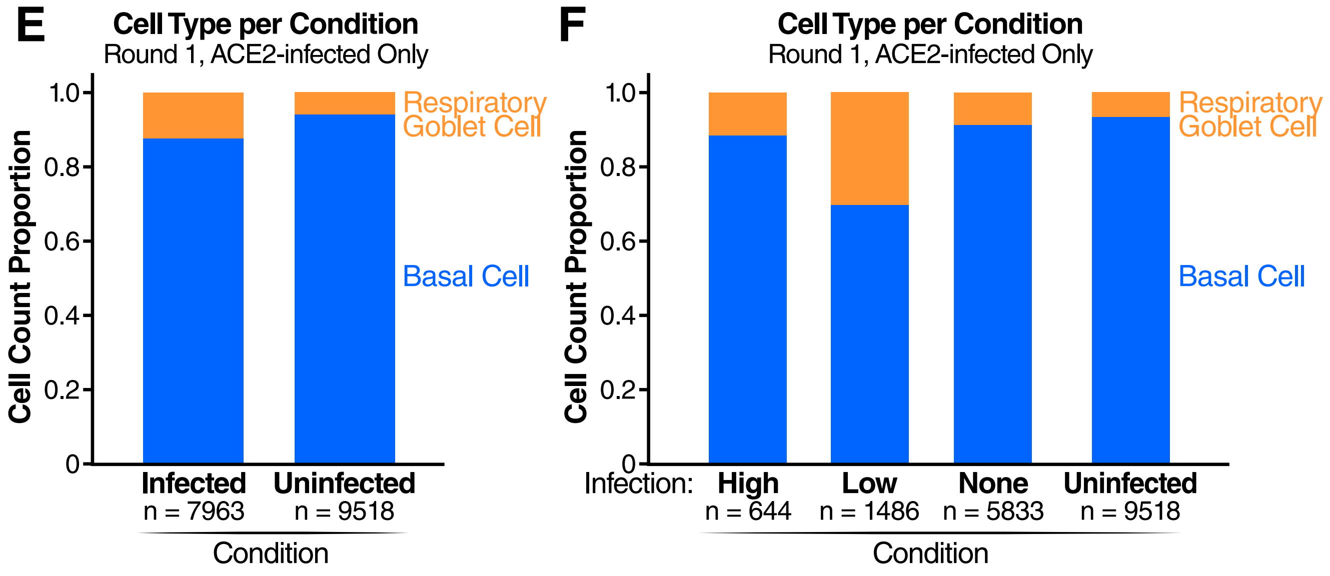


22.048 G C.Simoneau

**Supplementary Figure 2**

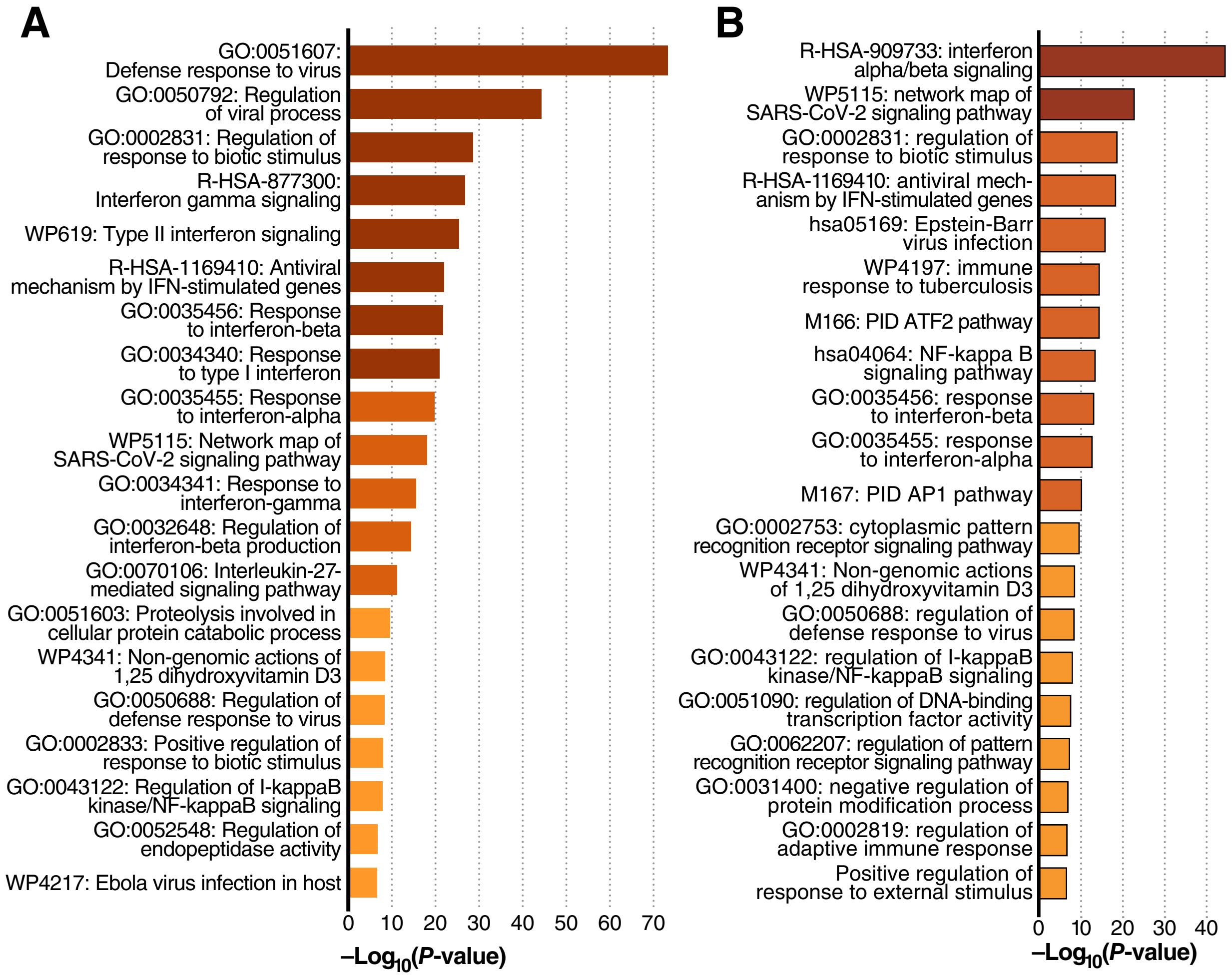


*22.048 H C.Simoneau*

### Supplementary Figure 3

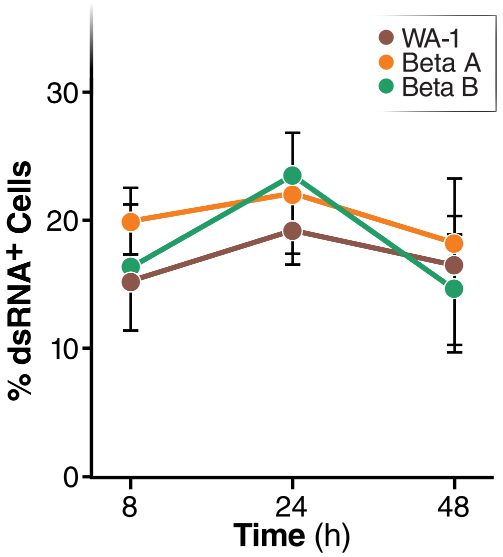

**Supplementary Figure 4**

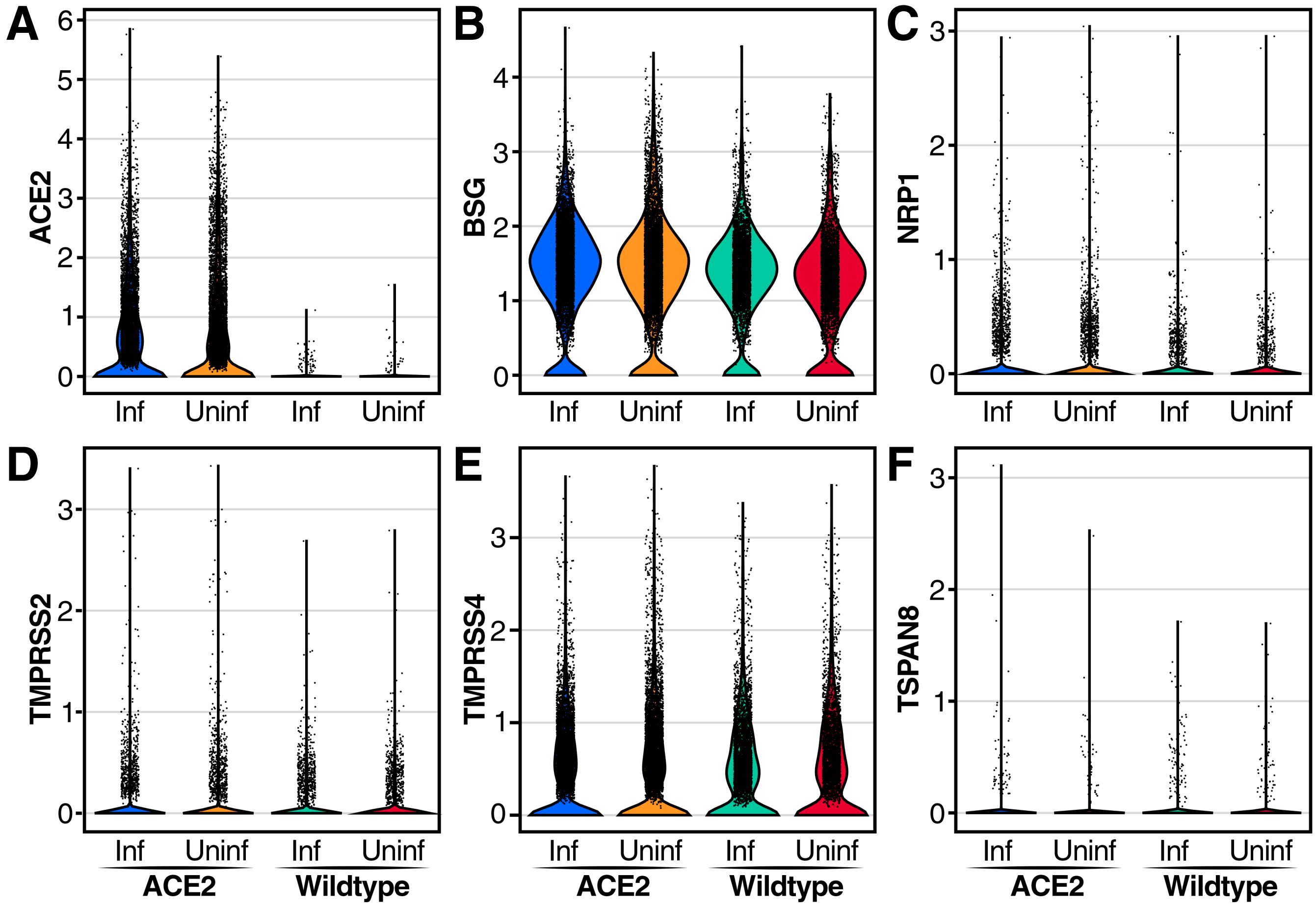


22.048 J C.Simoneau
